# DeepMP: a deep learning tool to detect DNA base modifications on Nanopore sequencing data

**DOI:** 10.1101/2021.06.28.450135

**Authors:** Jose Bonet, Mandi Chen, Marc Dabad, Simon Heath, Abel Gonzalez-Perez, Nuria Lopez-Bigas, Jens Lagergren

## Abstract

**Motivation:** DNA Methylation plays a key role in a variety of biological processes. Recently, Nanopore long-read sequencing has enabled direct detection of these modifications. As a consequence, a range of computational methods have been developed to exploit Nanopore data for methylation detection. However, current approaches rely on a human-defined threshold to detect the methylation status of a genomic position and are not optimized to detect sites methylated at low frequency. Furthermore, most methods employ either the Nanopore signals or the basecalling errors as the model input and do not take advantage of their combination.

**Results:** Here we present DeepMP, a convolutional neural network (CNN)-based model that takes information from Nanopore signals and basecalling errors to detect whether a given motif in a read is methylated or not. Besides, DeepMP introduces a threshold-free position modification calling model sensitive to sites methylated at low frequency across cells. We comprehensively benchmarked DeepMP against state-of-the-art methods on *E. coli*, human and pUC19 datasets. DeepMP outperforms current approaches at read-based and position-based methylation detection across sites methylated at different frequencies in the three datasets.

**Availability:** DeepMP is implemented and freely available under MIT license at github.com/pepebonet/DeepMP

**Contact:** —

## 1 Introduction

Chemical modifications of nucleotides are important epigenetic markers. These DNA modifications play a crucial role in regulating the expression of genes, and cellular responses to stimuli [3, 11, 25]. Among all the modifications, one of the most prevalent and widely-studied is DNA methylation, in particular at cytosines. Its relevance to fundamental biological processes such as embryonic development [18], aging [8], and diseases [9] has highlighted the importance of accurate genome-wide profiling techniques.

Both short and long-read sequencing technologies are exploited to identify methylated genomic sites. These approaches are associated with different experimental and computational limitations. For instance, the Bisulfite sequencing [20] with short reads suffers from limited conversion efficiency (Cytosine to Uracil). Also, the amplification bias and the uncertainty of mapping large repetitive regions add to the experimental complexity of its use for methylation detection. Long read-based technologies, such as Nanopore and PacBio sequencing approaches, have appeared more recently, and as a result, computational methods for DNA methylation detection are currently under development. Although PacBio Single-Molecule Real-Time (SMRT) approach can detect DNA methylation directly [6], low signal-to-noise ratios and coverage requirements have limited its application [5, 31]. Conversely, Nanopore sequencing overcomes these limitations as no amplification and prior enzymatic or chemical treatment steps are required, thus supporting the analysis of DNA molecules harboring modifications in their native state [13, 23, 24, 29]. Therefore, Nanopore sequencing has been recently established as the state-of-the-art long-read sequencing approach to detect DNA methylation.

In recent years, several models have contributed to the improvement of Nanopore’s methylation detection accuracy [16, 17, 19, 21, 22, 26, 27, 2]. The majority of these methods focus on directly analyzing the output signals of the Nanopore device and do not make use of the basecalling errors [17, 19, 21, 22, 26, 27]. These models have shown the ability to accurately call the methylated status for a target motif at the level of individual reads (read-based calling). However, the position-based calling (across all reads) relies on a human-defined threshold. An alternative approach to detect DNA methylations through Nanopore uses basecalling errors as model features [16]. Methods based solely on this approach can, thus far, only identify methylated genomic positions, rather than reveal the methylation status of every read covering the site. Therefore, these methods tend to underpredict sites methylated at low frequency across cells, i.e., sites at which most reads covering the base will be unmodified.

We hypothesized that the combination of basecalling errors and current signals could improve the detection of methylated cytosines in the DNA over the use of only one of them. Thus, we developed DeepMP, a deep learning model that exploits the errors and signals from the data. DeepMP includes a further innovation, a supervised Bayesian model to call the position-based methylation, which is –to our knowledge-unique. It comprises a statistical approach based on the information from individual reads covering a position that improves the detection of the methylation status of a genomic position. We show that DeepMP outperforms state-of-the-art methods DeepSignal [21], Nanopolish [26], Guppy [1] and Megalodon [2] in the tasks of detecting modified reads and positions methylated at varying frequencies across *E. coli*, human and pUC19 Nanopore long-read data.

## 2 Materials and methods

### 2.1 Datasets

Three datasets, K12 ER2925 (*E. coli*), NA12878 (human), and pUC19 (plasmid DNA) were employed to evaluate the performance and benchmark DeepMP. Table S1 summarizes all datasets and the number of reads available for each sample. The datasets employed in this study contain methylation on the cytosines (*E. coli* and human) and adenine (pUC19) residues. DeepMP was trained to distinguish unmethylated cytosines (C) from 5-Methylcytosines (5mC) at CpG motifs in the first two datasets and unmethylated adenines (A) from 6-methyladenines (6mA) at GATC motifs in the latter.

#### 2.1.1 CpG methylation data

Nanopore reads from *E. coli* (K12 ER2925) [26] were downloaded from the European Nucleotide Archive under accession number PRJEB13021. The dataset contains reads obtained from *E. coli* treated with M.SssI, which lead to methylation of ~ 95% of the CpGs (5mC methylated), and PCR-amplified (negative control), which are completely unmethylated. Although reads from both Nanopore R7.3 and R9 flow cells are included, R7.3 flow cells do not provide raw signals, and therefore only R9 reads were employed.

Human Nanopore reads from the NA12878 datasets were downloaded from the European Nucleotide Archive under accession number PRJEB23027 [10]. Reads are obtained from 5 sequencing studies (Norwich, UCSC, Bham, Notts, and UBC). However, only the Norwich subset is employed (Table S1). Datasets were basecalled by Guppy 4.4.2 (available to members of the Nanopore community at [1])

#### 2.1.2 Bisulfite sequencing data

Bisulfite sequencing results of the NA12878 datasets were downloaded from ENCODE (ENCFF835NTC) [4]. Both replicates available allow the labeling of high-quality methylated and unmethylated positions. We follow the approach of Liu et al. to characterize the methylation status of each position. If a cytosine in a given position in the reference genome (GRCh38) contained > 90% of methylations in both replicates, that position was considered to be modified and hence completely methylated. On the other hand, if a cytosine has 0% methylations in both replicates of bisulfite sequencing, the position was considered unmodified and thus completely unmethylated.

#### 2.1.3 6mA methylation data

Raw Nanopore reads from pUC19 were downloaded from NCBI under accession number SRR5219626. pUC19 plasmids were cloned in *E. coli* grown either in the presence (treated) or absence (untreated) of Dam methyltransferase (Table S1). The presence of this enzyme leads to the complete methylation adenines (m6A) in GATC motifs [22]. Sequences were also basecalled using Guppy 4.4.2 (available at: [1])

### 2.2 Benchmarking methylation detection with similar tools

We benchmarked DeepMP against four existing tools: Nanopolish [26], Megalodon [2], DeepSignal [21], and Guppy [1]. The different tools are explained in Section 1.1 of the Supplementary Methods.

### 2.3 Preprocessing and Feature Extraction

DeepMP consists of two different modules: a sequence module for handling the signals (Nanopore currents) and another module to process basecalling errors (Figure 1A, 1B). In the sequence feature extraction, the raw Nanopore reads are first basecalled using Guppy [1], and then re-squiggled by Tombo [27] to aligning the currents to the reference genome. Following Ni et al. [21] median normalization [27] was then applied to the raw signals. To obtain the error features, variants need to be called from fastq files by samtools [15] after the reference alignment using minimap2 [14]. The resulting files consist of the status of a position in a read (Match, Mismatch, Deletion, Insertion) and the quality of the call. This data is then processed to obtain the information of the presence or absence of mismatches, deletions, and insertions at the genomic position of the read.

**Figure 1:**
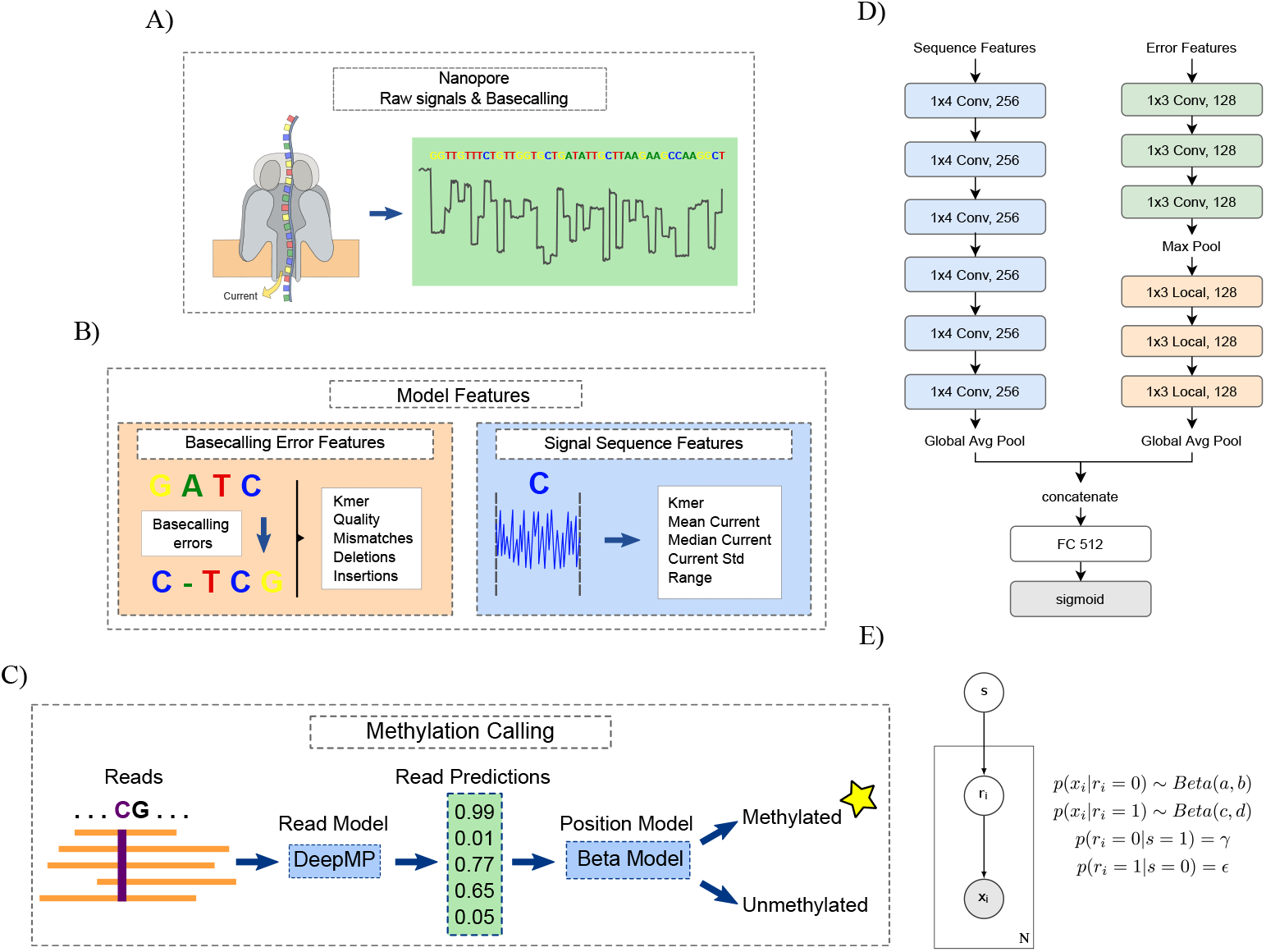
DeepMP model overview. **A)** Scheme of Nanopore technology, raw signals, and basecalling. **B)** Input features are divided into the basecalling error features (left) and the signal sequence features (right). Error features comprise the sequence information, read quality, mismatches, deletions, and insertions. Sequence features consist of the sequence data and the mean, median, std, and range of the base currents. **C)** Methylation calling pipeline. **D)** DeepMP architecture. The sequence module involves 6 1D convolutional layers with 256 1 × 4 filters. The error module comprises 3 1D convolutional layers and 3 locally connected layers both with 128 1 × 3 filters. Outputs are concatenated and inputted into a fully connected layer with 512 units. **E)** Probabilistic graphical model of the proposed Bayesian approach for position-based calling. The input data *x_i_* consists of *N* read predictions. *r_i_* is the read state, and *s* is the position state to be detected. Shaded nodes represent observed values, while the values of unshaded nodes are inferred.

Specifically, features for DeepMP are selected through an Incremental Feature Selection (IFS) strategy. We applied this strategy to a set of 11 features. Seven of them for the sequence module: mean, median, standard deviation, range, skewness, kurtosis, and the number of signals; and four for the error module: base quality, mismatches, deletions, and insertions. The features are ranked by the model performance (Figure S3, Figure S4) on one subset of the *E. coli* data and a subset of the human data (Figure S5). Both sample sets contain 100,000 labeled examples. The selection starts with the top 1 ranked feature (signal mean in *E. coli*) and iteratively adds features into the combination according to the rank until the end of the list is reached. If a decrease in performance is observed, the current feature will be excluded in the following run. A 5-fold cross-validation was employed to evaluate the performance of the selected features (Figure S4).

Based on this strategy, the features employed by DeepMP are the mean, median, standard deviation, and value range of the Nanopore signals for the sequence module and the read quality, deletions, insertions, and mismatches for the error module. After feature extraction for each target base, an *l*-length vector is constructed for every feature. Every vector contains the value of the nucleotide of interest and its *l* — 1 closest neighbors from both directions.

### 2.4 DeepMP Framework

Recent efforts to detect modifications liu2019detectionni2019deepsignal for Nanopore sequencing data have been exploring long short-term memory recurrent neural networks (LSTM RNNs), which are neural network models designed to learn long-term dependencies in sequential data. On a high level, our prediction task could be regarded as a many-to-one problem where the input of the model is a sequence and the output is a single value indicating the methylation state of the targeted base, which appears to naturally match the specialty of RNNs. However, this perspective overlooks some characteristics of the data: First, The inputs of the mentioned models are statistical measures of the signals instead of the raw signals, which have weaker time dependencies between the variables. Second, The base in interest locates at the middle of the input sequence rather than the end. In the mentioned methods, a bidirectional RNN first processes the sequence from one end to the other to output a representation for the whole sequence, then performs the same process for the reversed sequence. The representations obtained from this procedure are most sensitive to the input bases located at both ends of the sequencegoodfellow2016deep instead of the middle one. Besides, since these methods use a fixed-length sequence as the input to the networks, the flexibility of RNNs to adapt to various sequence lengths is not fully utilized in this particular problem setting. Additionally, the large amount of parameters in RNNs makes training the networks costly.

From a different aspect, one might consider the excerpted set of features to be a series of buckets containing information for each base in the sequence. The goal is to capture the interactions between the center bucket and its neighbors, as well as the interactions among them. Concerning the spatial correlation in these buckets, we propose to employ convolutional neural networks (CNNs). By virtue of sparse connections between units and the parameter sharing in the convolutional layers, CNNs are memory efficient and easy to parallelize, therefore, less expensive to train compared to RNNs. CNNs have been successfully applied to numerous tasks such as classification, detection, segmentation in various study fields including computer vision, natural language processing, drug discovery, etc. In our DeepMP model, there are two CNN modules (Figure 1D): the sequence module and the base-calling error module. They are designed to process two different sets of features extracted from Nanopore sequencing data.

#### 2.4.1 Sequence Module

The four *l*-length vectors generated by sequence feature extraction are stacked with the one-hot embedded nucleotide sequence to form the model input vector of size *l* × 9. The input vector is then given to the sequence CNN module, which is composed of 6 1D convolutional layers with 256 1 × 4 filters. The stride for convolution is fixed to 1 base. The convolution function computes the nth element *Z* on the feature map by

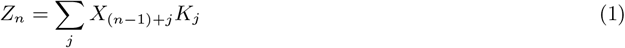

where *X* is the input to the layer, *K_j_* is the *j*th element in kernel tensor.

Batch normalization (BN) is applied to each convolutional layer right before a ReLU activation function. The sequence module ends with a global average pooling layer.

#### 2.4.2 Basecalling Error Module

With the same treatment as to the sequence feature, the *l*-length vectors are stacked with the one-hot embedded nucleotide sequence, resulting in the final *l* × 9 shaped input for the error module.

The error module contains two types of layers: 1D convolutional layers and locally connected layers le-cun1989generalization. The convolutional layers compress the features into a compacted representation whereas the locally connected layers detect the neighborhood information in the sequence. A locally connected layer learns a set of filters separately for every location, naturally, it captures spatial characteristic information of the features. Since the signals from one base in the DNA sequence is highly correlated with its close neighboring bases, learning the spatial patterns enhances the model’s ability to discriminate between different modification states. The input vector is fed into a three-layer convolutional neural network followed by a max-pooling layer, and then a three-layer locally connected network, finally a global average pooling layer. Both of the convolutional layers and locally connected layers have 128 1 × 3 filters.

#### 2.4.3 Model Outputs

The outputs from the sequence module and the error module are concatenated and inputted into a fully connected layer with 512 units. Later on, the last fully connected layer with a sigmoid activation function outputs the final prediction *ŷ* ∈ [0, 1] for the central base. Additionally, the two modules can be independent models by themselves.

#### 2.4.4 Training DeepMP

During the training process, the model learns to minimize the loss computed by the binary cross-entropy loss function

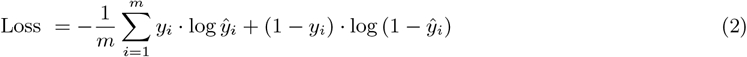

where *y* is the label, *ŷ* is the model output, and *m* is the mini-batch size.

We trained the model using an Adam optimizer with a learning rate of 0.00125 and a mini-batch size of 512, early stopping was applied to prevent overfitting. In the default settings, we set the sequence length *l* to be 17. The full implementation is in Python 3 with Tensorflow 2.

In the experiments, DeepMP was trained for 10 to 20 epochs depending on the size of the dataset. On *E. coli* we performed a 5-fold Cross-validation following the strategy proposed by Liu et al [17]. The *E. coli* genome was split into five sections ([0, 1000000], [1000000, 2000000], [2000000, 3000000], [3000000, 4000000], [4000000, 4700000]) (Table S2). Then, models were trained on four sets and test on the remaining one. For the human dataset chromosome 1 was kept for testing, while other regions were used for training and validation. For the pUC19 plasmid DNA data, reads were split into a 90% training set, a 5% validation set, and a 5% test set. In all datasets, training, test, and validation sets contained 50/50 positive and negative samples.

#### 2.4.5 Position Modification calling

Individual read-based calls mapped to a certain position in the genome need to be gathered to predict the methylation status of that position. Particularly, we introduce a Bayesian approach to accomplish the position modification calling (Figure 1E). Once the neural networks are tested on labeled data, read-based predictions for each label group can be obtained. These values are subsequently used to infer the parameters of the underlying distribution for each group. By incorporating the read-based predictions and the ground truth labels, the proposed Bayesian approach provides more precise calls for the genomic positions.

The Bayesian model is based on the following: let **x** = {*x*_1_, *x*_2_,…, *x_N_* } ∈ Ω*^N^* to be the prediction of *N* reads for one position. Assuming different read calls are independent and identically distributed, the probability of observing **x** given the position state *s* can be computed by

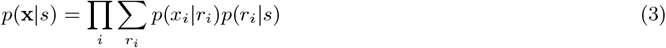

where *r_i_* indicates if the *i*-th read mapped to the position is modified, *r_i_* ∈ {0, 1}. The graphical model is shown in Figure 1E.

According to Bayes’ rule, given a prior distribution *p*(*s*) the posterior probability of s is in proportion to *p*(**x**|*s*)

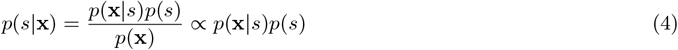

To infer the position state we need to compute *p*(**x**|*s* = 0) along with *p*(**x**|*s* = 1) and compare these two likelihoods. Notice that given position state *s*, the random variable *r_i_* follows a Bernoulli distribution. Therefore, we model *p*(*r_i_*|*s*) with two Bernoulli distributions parameterized by *ε* and *γ*:

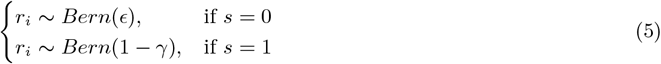

then model *p*(*x_i_*|*r_i_*) with beta distribution parameterized by *a, b, c, d* ~ {*z* ∈ *R* | *z* ≥ 0}

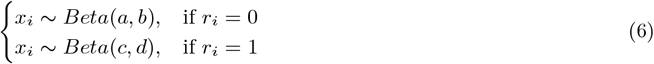

with equations 3-6 we arrive at the final expression:

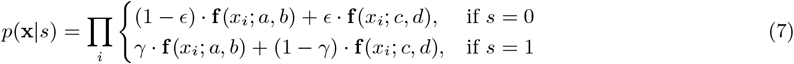

where **f** is the probability density function of beta distribution.

We now discuss how to choose parameters *γ, ε* and *a, b, c, d*. The probability of having a negative read state on a positive position is relatively high, whereas having a positive read state on a negative position is rather a rare event. Therefore, *γ* should be set to a value close to 1 while *ε* should be close to 0. In the experiment, we fix *γ* = 0.83 and *ε* = 0.05. The parameters for the beta distribution are estimated from the read-based predictions on a independent labeled dataset. The predictions are divided into two groups according to the label of the sample, where we estimate the sample mean *μ* and variance for each group. The shape parameters *α* and *β* for beta distribution can be approximated by

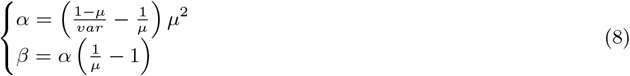

finally, we obtain two sets of *α* and *β*: *a, b* for label 0 group and *c, d* for label 1 group.

#### 2.4.6 Performance Evaluation

We quantify the performance of the models by four common measures: accuracy, precision, recall, and F-score.

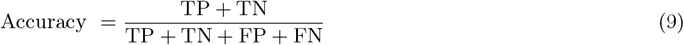

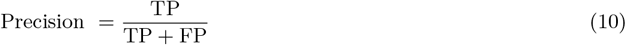

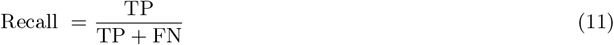

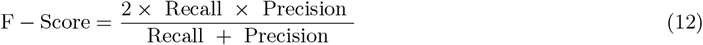

where TP, TN, FP, FN, are abbreviations for true positives, true negatives, false positives, and false negatives, correspondingly. The accuracy gives an overview of the quality of the predictions, while precision shows the correct predictions in the positive calls, and recall is the percentage of correct predictions in all positive samples. Finally, the F-score is the harmonic mean of precision and recall which shows the balance between the two.

## 3 Results

DeepMP takes as input two types of information from Nanopore sequencing data (Figure 1A), basecalling errors and raw current signals (Figure 1B). Features from these two types of information are fed into a CNN-based module to identify modified sites in individual reads (Figure 1D). These read level predictions are then integrated by the position-based calling method described in Section 2.4.5 (Figure 1E), to identify methylated genomic sites (Figure 1C).

We assessed the performance of DeepMP in the task of detecting methylated sites on three different datasets, *E. coli*, human and pUC19. The *E. coli* dataset has been widely used for model training and benchmarking. It consists of a negative control with 0% methylation (PCR amplified) and a positive one treated with a methyltransferase (M.SssI) that converts 5C to 5mC with a ~ 95% efficiency [26] (Section 2.1.1). Regarding the human dataset, the available bisulfite sequencing allowed the labeling of high-quality methylated and unmethylated positions (Section 2.1.2). Moreover, the pUC19 dataset opens up the possibility to evaluate DeepMP in the identification of a different modification (6mA) within GATC motifs. These three datasets were used to benchmark DeepMP against Megalodon [2], Guppy [1], DeepSignal [21] and Nanopolish [26].

### 3.1 5mC detection performance on *E. coli* data

DeepMP and DeepSignal were trained on a mixture of methylated and unmethylated reads as described in Section 2.4.4. For the other methods, we used the available pre-trained models and the procedures explained in Section 2.2. All methods were tested on the same set of the data.

#### 3.1.1 Prediction accuracy and studies at read level

To determine whether a genomic position is methylated, the first step is to detect the methylation status of each read covering it (Figure 1C). Thus, an independent set of reads, as described in Section 2.4.4 is selected to evaluate all three models. DeepMP showed the best Area under the ROC (AUC) among all the benchmarked methods (DeepMP: 0.988, Megalodon: 0.981, DeepSignal: 0.974, Guppy: 0.924, Nanopolish: 0.878) (Figure 2A). While Guppy and Megalodon presented the highest precision values (Guppy: 0.9957, Megalodon: 0.9906), DeepMP exhibited the highest overall accuracy (0.9397), recall (0.9347), and f-score (0.9398) (Figure 2B). These results were consistent throughout the 5-fold cross-validation (Figure 2C; Table S2).

**Figure 2:**
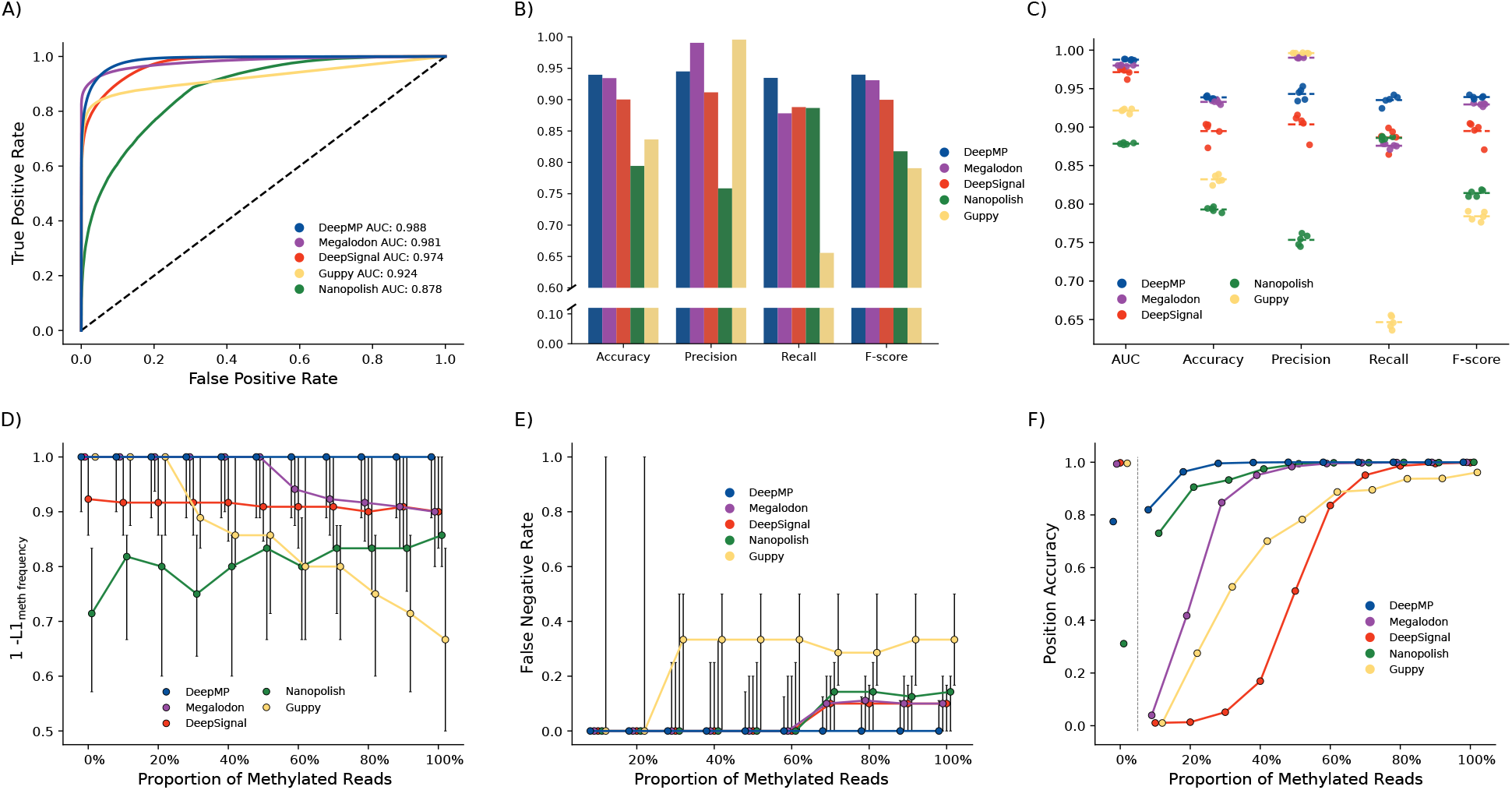
Performance of DeepMP, Megalodon, Guppy, DeepSignal, and Nanopolish on *E. coli* dataset. **A)** Receiver operating characteristic (ROC) curve showing the false positive rate (x-axis) vs. the true positive rate (y-axis) for the read predictions on a mixture of methylated and unmethylated reads in a single cross validation fold. **B)** Accuracy measurements for the compared models in the cross validation fold in A. **C)** 5-fold cross validation accuracy and AUC results for mixtures of methylated and unmethylated reads **D)** Accuracy to determine the methylation frequency of a sample measured by 1 - *L*1_meth frequency_ (y-axis). 1 - *L*1_meth frequency_ is evaluated on eleven datasets comprising different levels of methylated reads (0-100%) (x-axis). **E)** False negative rate (y-axis) evaluated on the same eleven datasets as in D. **F)** Position calling accuracy for each of the partially methylated datasets at different thresholds (DeepMP: Bayesian approach; DeepSignal: mean read predictions estimate; Megalodon, Guppy and Nanopolish: 20 % threshold)

These read predictions come from a mixture of two samples that are completely methylated or unmethylated. However, in a natural population of cells, one particular base may be methylated across some, but not all of them. For instance, 6mA across 12 different genomic sites is found at levels ranging from 7 to 69% in yeast samples [7]. To benchmark DeepMP in a real-life scenario, we thus generated 11 synthetic datasets featuring mixtures of methylated and unmethylated reads at different proportions. The accuracy was measured as 1 — *L*1_meth frequency_. The *L*1_meth frequency_ distance is defined as the absolute value of the difference between the true methylated frequency of the position and that estimated by the model. This measurement takes into account FP and FN at read-level, avoiding biased estimates of the methylated frequency. Figure 2D shows the improved performance of DeepMP for inferring the true methylated frequency of the sample. The accuracy of DeepMP was the highest among the methods at high proportion of methylated reads. At low proportions of methylated reads, Megalodon and Guppy showed a comparable performance. These results are in accordance with the FN and FP levels shown in Figures 2E and S1A. DeepMP presents a lower number of FN across all proportion of methylated reads, and the levels of FP are consistent across synthetic datasets for all methods except Nanopolish.

One of the main innovations of DeepMP consists in combining a base-calling error module and a sequence signal module. To assess how much this innovation actually contributes to the observed gain in performance, DeepMP trained on the sequence module only (DeepMP Seq) was also included in the comparison (Figure S1C, S1D). DeepMP Seq shows a decreased performance compared to DeepMP (DeepMP F-score: 0.9398; DeepMP Seq F-score: 0.889) (Table S2), which highlights the importance of taking into account the features derived from basecalling errors for the *E. coli* dataset.

#### 3.1.2 Prediction accuracy on genomic sites

The ultimate goal of methods that identify DNA methylation is to be correctly determine the frequency of methylation of a genomic site across cells. This is done through the identified read level methylation sites of reads overlapping the genomic position under analysis. One strategy to do so is to decide a particular percentage of methylated reads as the threshold [17], i.e., a 20% threshold will detect as methylated any position presenting more than 20% of methylated overlapping reads. Another approach is to get a mean estimate of the predictions of the *n* reads overlapping a particular position [21].

Although such approaches yield acceptable results on datasets with sites methylated at high-frequency [17, 21], their accuracy usually drops at sites methylated at lower levels (10-30%) (Figure 2F) (a 20 % threshold was applied to all models but DeepSignal and DeepMP). To solve this problem, we propose to apply a Bayesian approach (DeepMP Figure 2F; Figure 1E), which utilizes the statistical features of the read predictions to compute the likelihood of the different states for the genomic position (Section 2.4.5). As a result, DeepMP shows the highest accuracy (*>* 80% of the positions) when the dataset contains, on average, sites methylated in 10% of the reads (Figure 2F), while robustly calling ~ 80% of the positions when the dataset is completely unmethylated. This trend is conserved when comparing DeepMP using different thresholds (10%, 20% and 50% Threshold) (Figure S1B).

### 3.2 5mC detection performance on human data

DeepMP and DeepSignal were trained on reads containing methylated and unmethylated cytosines (determined by bisulfite sequencing) and tested on the same type of data drawn from reads overlapping human chromosome 1 in the Norwich subset of the human data (Table S1). For Megalodon, Guppy, and Nanopolish, we used available pre-trained models as explained in Section 2.2 and we tested them on the same set of data.

#### 3.2.1 Prediction accuracy at read level

Similar to *E. coli* dataset, DeepMP, Megalodon, Guppy, DeepSignal, and Nanopolish were evaluated on their read prediction performance at read level (Figure 3A, B; Table 1). DeepMP achieves the better ROC AUC scores (DeepMP: 0.967; Megalodon: 0.9394; Guppy: 0.8602; DeepSignal: 0.9629; Nanopolish: 0.9284) and F-scores (DeepMP: 0.9324; Megalodon: 0.8968; Guppy: 0.7787; DeepSignal: 0.9255; Nanopolish: 0.9236).

**Figure 3:**
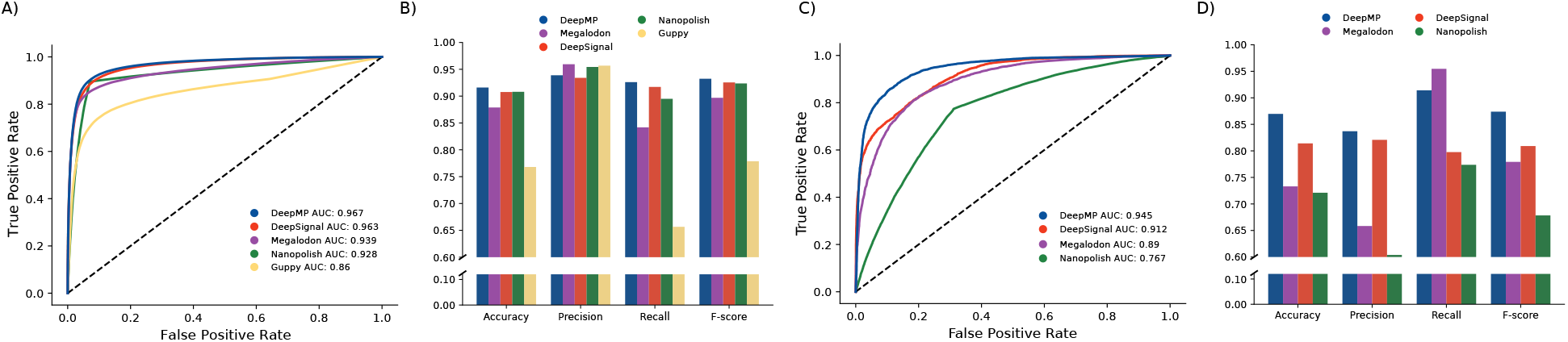
Performance of DeepMP, Megalodon, Guppy, DeepSignal and Nanopolish on the Norwich subset of the human dataset and of DeepMP, Megaldon, DeepSignal and Nanopolish on the pUC19 plasmid. **A, C)** Receiver operating characteristic (ROC) curve showing the false positive rate (x-axis) vs. the true positive rate (y-axis) for the read predictions on a mixture of methylated and unmethylated reads on the human dataset (A) and pUC19 (C) respectively. **B)** Accuracy measurements for the models on the human dataset. **D)** Accuracy measurements for the models on the pUC19 plasmid

**Table 1:** Performance summary at read level of DeepMP, Nanopolish, DeepSignal, Guppy and Megalodon on *E. coli*, human and pUC19 datasets. Accuracy measurements for *E. coli* represent the average value of the 5-fold cross validation

| Dataset | Test | Method | Accuracy | Precision | Recall | F-Score |
| --- | --- | --- | --- | --- | --- | --- |
| *E. coli* | Read level | Nanopolish | 0.7929 | 0.7535 | 0.8859 | 0.8143 |
|  |  | DeepSignal | 0.8948 | 0.9035 | 0.8865 | 0.8949 |
|  |  | Guppy | 0.8322 | **0.9961** | 0.6466 | 0.7841 |
|  |  | Megalodon | 0.9327 | 0.9901 | 0.8757 | 0.9294 |
|  |  | DeepMP | **0.9386** | 0.9431 | **0.9352** | **0.9391** |
| H. sapiens | Read level | Nanopolish | 0.9080 | 0.9543 | 0.8949 | 0.9236 |
|  |  | DeepSignal | 0.9076 | 0.9340 | 0.9191 | 0.9255 |
|  |  | Guppy | 0.7680 | 0.9568 | 0.6565 | 0.7787 |
|  |  | Megalodon | 0.8788 | **0.9594** | 0.8418 | 0.8968 |
|  |  | DeepMP | **0.9160** | 0.9388 | **0.9260** | **0.9324** |
| pUC19 plasmid | Read level | Nanopolish | 0.7211 | 0.6038 | 0.7737 | 0.6783 |
|  |  | DeepSignal | 0.8141 | 0.8207 | 0.7976 | 0.8090 |
|  |  | Megalodon | 0.7730 | 0.6583 | **0.9547** | 0.7792 |
|  |  | DeepMP | **0.8697** | **0.8370** | 0.9141 | **0.8738** |

When comparing DeepMP Seq (DeepMP trained only using the sequence module) against DeepMP, Megalodon, DeepSignal, Guppy, and Nanopolish, it outperforms all methods but DeepMP (DeepMP Seq AUC: 0.966; DeepMP Seq F-score: 0.9315) (Figure S2A, S2B; Table S3). These results differ from the *E. coli* dataset where DeepMP Seq is behind DeepSignal in performance measurements.

### 3.3 6mA detection performance on pUC19 plasmid DNA data

To evaluate the ability of DeepMP to detect a different type of methylation (6mA), we benchmarked DeepMP on the pUC19 dataset. The read-based strategy was employed to train DeepMP and DeepSignal on a mixture of methylated (6mA) and unmethylated (A) reads as described in Section 2.4.4. As the model selected for Guppy does not specifically detect modifications in all contexts (Section 2.2), it was discarded. Nevertheless, Megalodon, considered (and supported by our previous results) the state-of-the-art Nanopore method for methylation detection ([30, 2]), rather than Guppy, was run with the available model for all contexts as described in Section 2.2. Nanopolish was run to detect modifications in *dam* sites (DNA adenine methylase). Considering the number of reads discarded by the selective cutoff, we tested Nanopolish on all the extracted examples. The rest of the models used the same set of data for testing.

DeepMP significantly outperforms DeepSignal, Megalodon, and Nanopolish in the detection of 6mA within GATC motifs (Table 1). In figures 3C and 3D, DeepMP also shows an improvement of performance in the binary classification task and the related accuracy measurements. Furthermore, DeepMP Seq alone was also able to outperform any other method but DeepMP in the detection of modified GATC motifs (Figure S2C, S2D; Table S3).

## 4 Discussion

Nanopore sequencing, coupled with the computational methods that exploit its output to detect base modifications, has opened up the possibility of directly identifying epigenetic modifications of DNA nucleotides. The advent of this technology has the potential to supersede methodologies like bisulfite sequencing, which is currently the gold standard to detect DNA methylation. For this to be accomplished, the accuracy computational methods of detection need to be improved to correctly identify the methylation level at individual genomic positions [7]. This study proposes a neural network solution, DeepMP, which utilizes both electric currents and basecalling errors from Nanopore sequencing to deliver high accurate methylation detection.

We showed that DeepMP outperforms state-of-the-art methods Megalodon, Guppy, DeepSignal, and Nanopolish on *E. coli*, human and pUC19 datasets. DeepMP favorably compared in the proposed benchmark when inferring the true methylated frequency of partially methylated *E. coli* datasets (0%, 10%, …, 100%), consistently presenting lower false postive and negative rates. Furthermore, the analysis of the methods’ ability to call the position methylation status displayed the limitation of introducing an arbitrary, human-defined threshold. A 20% threshold could not properly capture a position’s true methylation status in datasets with low methylation levels (¡30% methylation). Importantly, employing DeepMP’s novel Bayesian approach corrected this problem while being robust on completely methylated samples.

Intriguingly, in the case of the human dataset, the features based on basecalling errors do not appear to provide the same level of information as in the *E. coli* dataset. This finding indicates that the informational value of basecalling errors varies in different types of data, i.e., the error features may be dataset-specific. A hypothesis would be that the M.SssI treatment in *E.coli* samples generates a characteristic error pattern that is not present when mutations occur naturally (human dataset). Consequently, to obtain a more generalizable model across species, one might consider employing DeepMP without the basecalling errors module.

Moreover, we demonstrated that DeepMP detects not only 5mC but also 6mA more accurately than state-of-the-art methods. Given the low number of reads, DeepMP achieved a significant separation compared to Megalodon, DeepSignal, and Nanopolish. This fact highlights the versatility of DeepMP across different types of nucleotide modifications (including at certain sequence motifs) and sequencing coverage. Nevertheless, it shares one of the current limitations of most supervised learning models, namely, the inability to detect out-of-sample modifications. That is, to detect methylated bases absent in the training set. One way to overcome could be to use unsupervised or one-shot learning methods ([12, 28]. This technical advancement would allow the exploration of a different set of interesting problems regarding the identification of modified bases using Nanopore.

In summary, DeepMP accurately detects methylated sites both, in individual reads and at genomic positions, at different levels of methylation of the sample, as real-world biological samples. Besides, the proposed Bayesian approach could apply to related problems to substitute a human-defined threshold, especially when labeled data is available. In addition, we posit that DeepMP could also be employed to generate genome-wide maps of different DNA base lesions from Nanopore sequencing data. This could largely reduce experimental hurdles to achieve this objective. As Nanopore technology continues to advance and the sequencing costs are reduced, the amount of data will exponentially increase. Accurate and efficient methods as DeepMP will be key to exploit this sequencing data in scenarios with limited computational resources.

## Acknowledgements

IRB Barcelona is a recipient of a Severo Ochoa Centre of Excellence Award from the Spanish Ministry of Economy and Competitiveness (MINECO; Government of Spain) and is supported by CERCA (Generalitat de Catalunya).

