## Supplementary Material for "DeepMP: a deep learning tool to detect DNA base modifications on Nanopore sequencing data"

### 1 Supplementary Methods

#### 1.1 Benchmarking methylation detection with similar tools

We benchmarked DeepMP against four existing tools: Nanopolish [13], Megalodon [4], DeepSignal [11], and Guppy [3].

Nanopolish (v0.13.2) [13] assigns a log-likelihood ratio to each CpG site or group of nearby CpG sites that share the same methylation level. A positive log-likelihood ratio value indicates evidence of methylation. Groups of CpGs were separated to include all the predictions per read and site. Additionally, a site was considered to be methylated if the log-likelihood was above 2.0 and unmethylated if it was lower than -2.0 [2]. Finally, log-likelihood values were transformed into probabilities for the benchmarking. Methylation calling through Nanopolish was performed following the steps presented by Simpson et al. in [5].

DeepSignal (v0.1.7) [11] is an RNN-based model to detect the methylation state of a given motif. It renders information about the methylated status of the motif per read and site. If the probability of methylation was above 0.5, the position in the read was considered to be methylated. Otherwise, called unmethylated. Models for DeepSignal were trained and tested in the same sets as DeepMP.

Guppy (v4.4.2) [3] directly basecalls 5mC at CpG sites as a fifth base along the four canonical DNA bases. The configuration file `dna_r9.4.1_450bps_modbases_dam-dcm-cpg_hac.cfg` was used as Guppy's modified basecalling model. The reads produced incorporate the information supporting the modifications at a given genomic position. We then used the scripts provided at [1] to convert the guppy methylation calls from fast5 files to reference anchored files. As in nanopolish, resulting log-likelihood values were transformed into probabilities for the benchmarking.

Megalodon (v2.3.2) [4] uses Guppy to re-basecall the fast5 reads to then identify methylated motifs by aligning the basecalling output to the reference. It assigns a score for the candidate modified base and performs a calibration when converting the raw scores to estimated empirical probabilities. We employed the most recent basecalling models in Rerio [6] to detect CpG (`res_dna_r941_min_modbases_5mC_CpG_v001.cfg`) as well as methylation in GATC motifs (`res_dna_r941_min_modbases-all-context_v001 .cfg`) with Megalodon. Megalodon produces per-read modified base log probability and canonical base log probability at each mapped site of the motif. For the benchmarking analysis per site and read, we transformed the resulting log-likelihoods into probabilities.

Additional tools were not included for the following reasons. We decided not to include DeepMod [9] because DeepSignal, with a model with similar architecture, has been proven to outperform it [15]. Similarly, SignalAlign [12] has shown lower performance than DeepSignal [11]. Besides, the model has not been updated in recent years. We did not consider mCaller [10], because only models trained for 6mA and not for 5mC were available. Finally, Tombo [14] was not included in the analysis as it is not considered the state-of-the-art method to detect methylations by their developers, Oxford Nanopore Technologies. Furthermore, recent studies have shown reduced performance when compared against the methods employed in the presented benchmarking [15]. On the other hand, Epinano [8], a method to detect modifications based on the basecalling error information, was discarded due to its inability to call methylated sites per read.

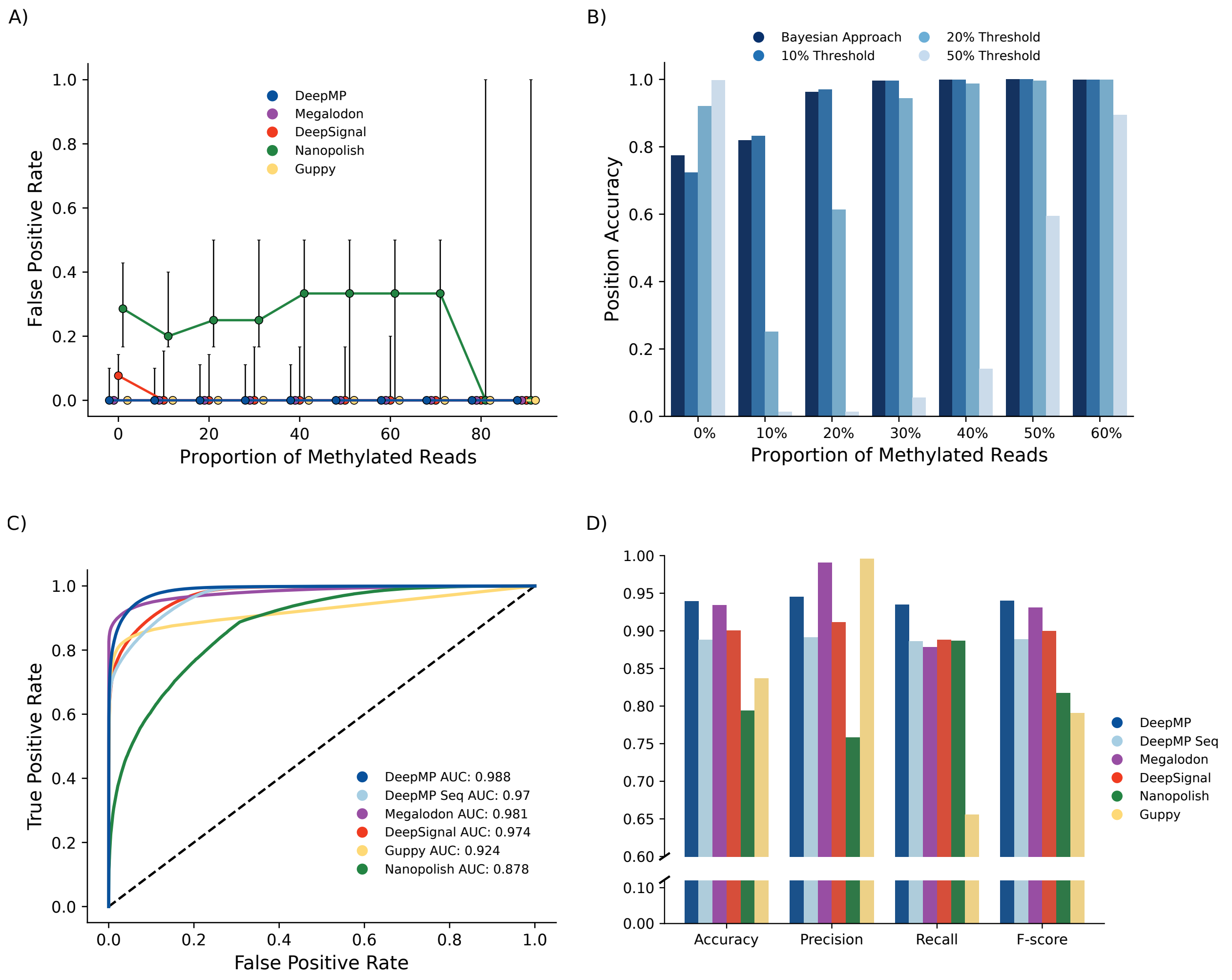


Figure S1: **A)** False positive rate (y-axis) evaluated on eleven datasets comprising different levels of methylation (0-100%) (x-axis) on DeepMP, Megalodon, DeepSignal, Guppy and Nanopolish. **B)** Comparison of the position accuracy of DeepMP at different thresholds and proportion of methylated reads against the position-based calling Bayesian approach. **C)** Receiver operating characteristic (ROC) curve showing the false positive rate (x-axis) vs. the true positive rate (y-axis) for the read predictions on a mixture of methylated and unmethylated reads in a single cross validation fold. **D)** Accuracy measurements for the models on the cross validation fold presented in C.

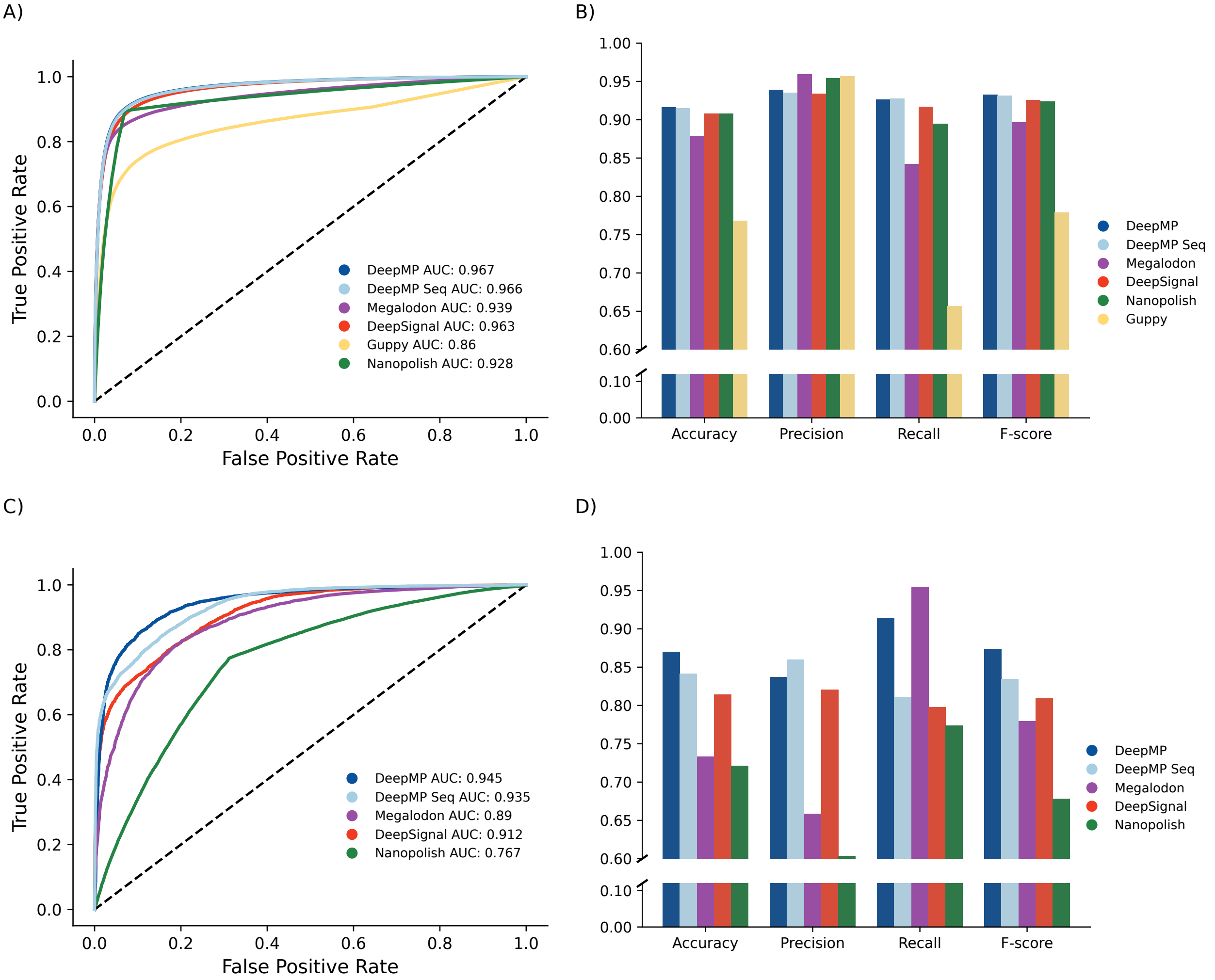


Figure S2: Performance of DeepMP, DeepMP Seq, Megalodon, Guppy, DeepSignal and Nanopolish on the Norwich subset of the human dataset and of DeepMP, DeepMP Seq, Megaldon, DeepSignal and Nanopolish on the pUC19 plasmid. **A, C)** Receiver operating characteristic (ROC) curve showing the false positive rate (x-axis) vs. the true positive rate (y-axis) for the read predictions on a mixture of methylated and unmethylated reads on the human dataset (A) and pUC19 (C) respectively. **B)** Accuracy measurements for the models on the human dataset. **D)** Accuracy measurements for the models on the pUC19 plasmid

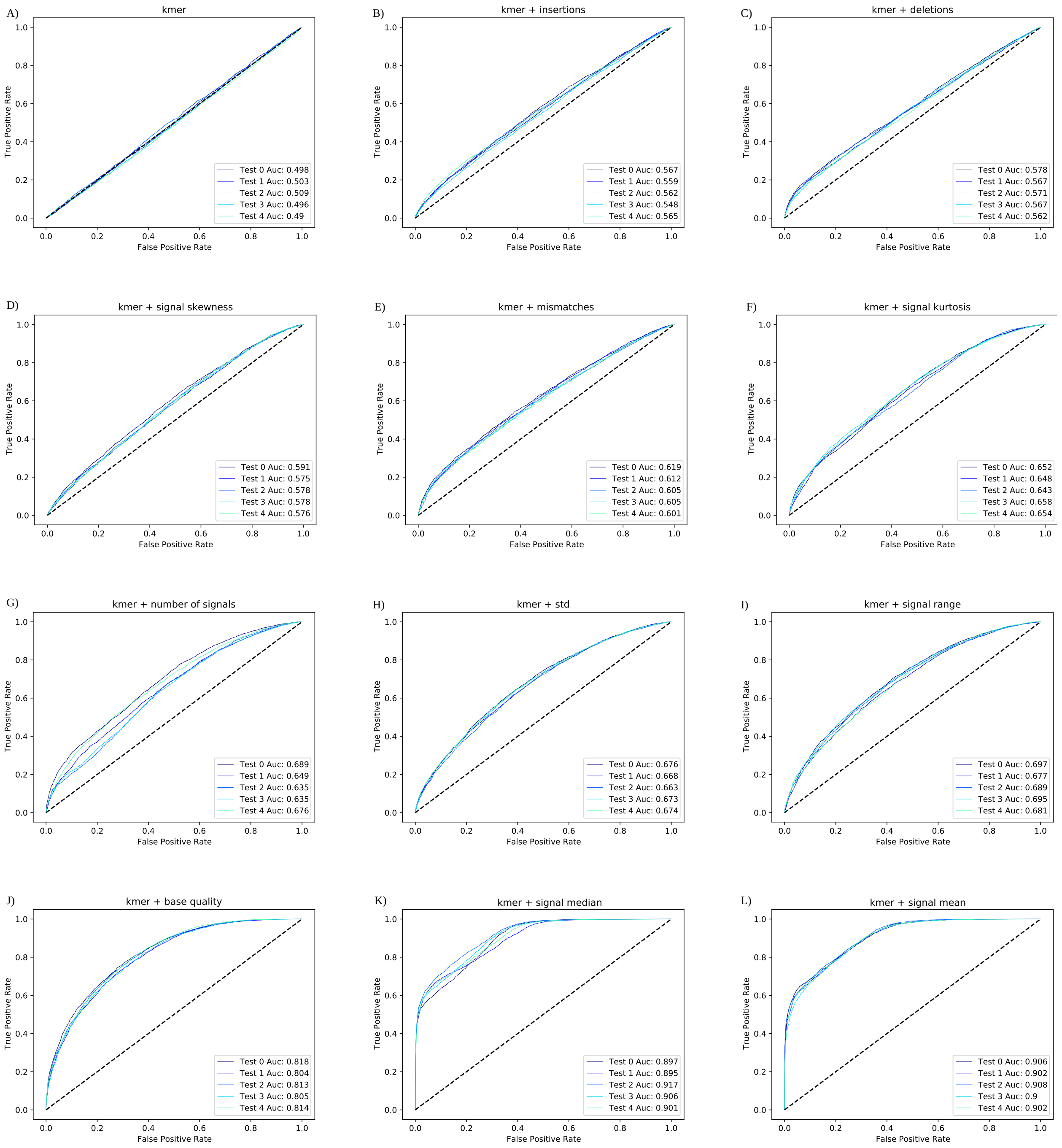


Figure S3: Single feature performance on a subset of the *E. coli* dataset. Kmer serves as a baseline feature and it is combined with all other individual features. Performance on **A)** kmer alone, **B)** insertions, **C)** deletions , **D)** signal skewness, **E)** mismatches, **F)** signal kurtosis, **G)** number of signals, **H)** signal std, **I)** signal range, **J)** base quality, **K)** signal median and **L)** signal mean. **A-L)** Receiver operating characteristic (ROC) curve showing the false positive rate (x-axis) vs. the true positive rate (y-axis) for the read predictions on a mixture of methylated and unmethylated reads. This analysis was conducted through a 5-fold cross-validation (Test 0-4) using 100,000 labeled examples of *E. coli* data.

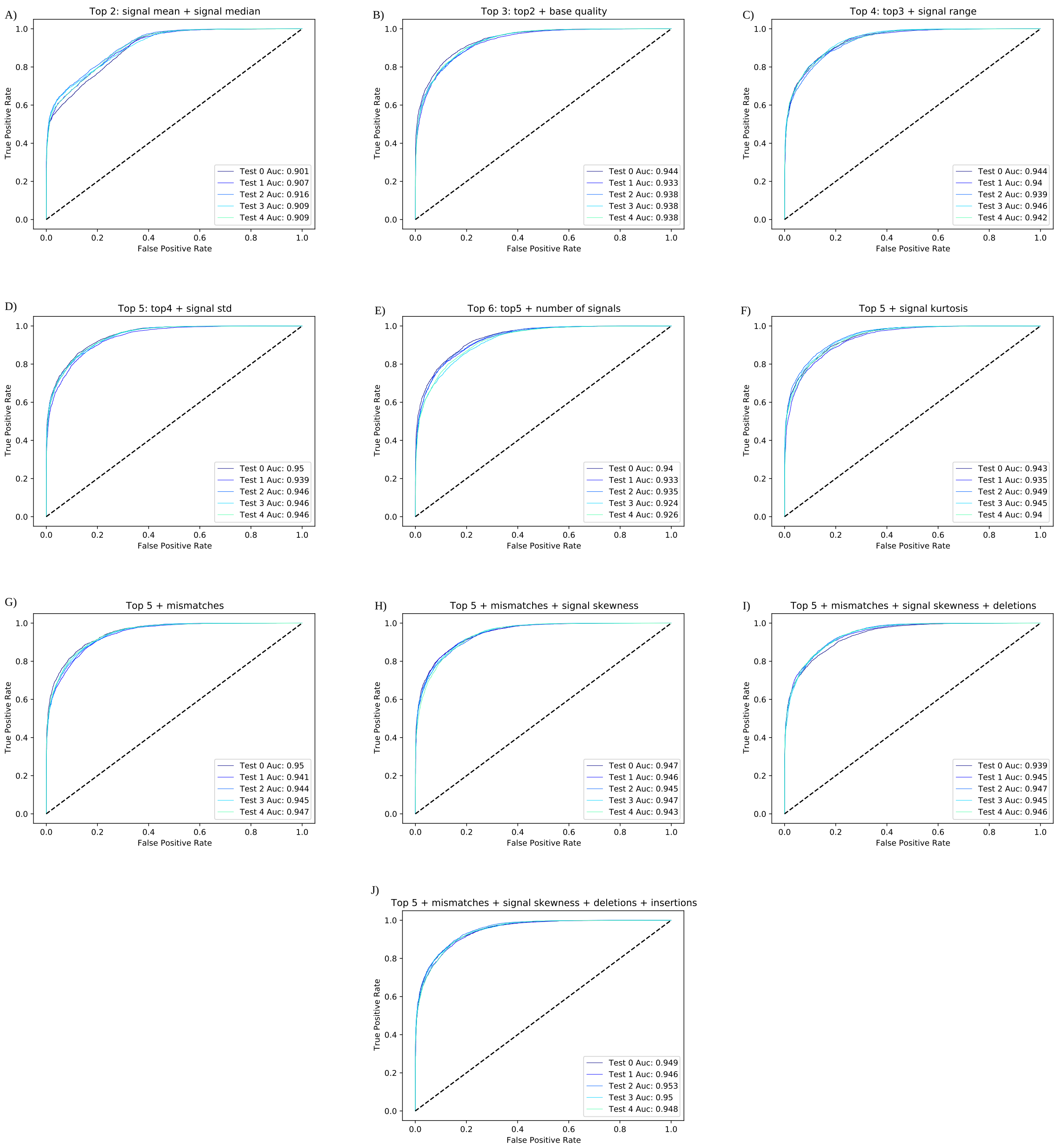


Figure S4: Incremental Feature Selection (IFS) combinations performance on a subset of the *E. coli* dataset. The rank of the 11 features by average AUC is 1. signal mean 2. signal median 3. base quality 4. signal range 5. signal std 6. number of signals 7. signal kurtosis 8. mismatches 9. signal skewness 10. deletions 11. insertions. The IFS strategy starts with **A)** signal mean and signal median and it adds one feature at a time to the combination for the model performance evaluation. **B, C, D, H, J)** The new feature is kept if it improves the performance. As an exception, Signal kurtosis is discarded due to its low AUC score shown in human data (S5). **E, G, I)** Otherwise, if the performance decreases, the new feature is discarded. **A-J)** Receiver operating characteristic (ROC) curve showing the false positive rate (x-axis) vs. the true positive rate (y-axis) for the read predictions on a mixture of methylated and unmethylated reads. This analysis was conducted through a 5-fold cross-validation (Test 0-4) using 100,000 labeled examples of *E. coli* data.

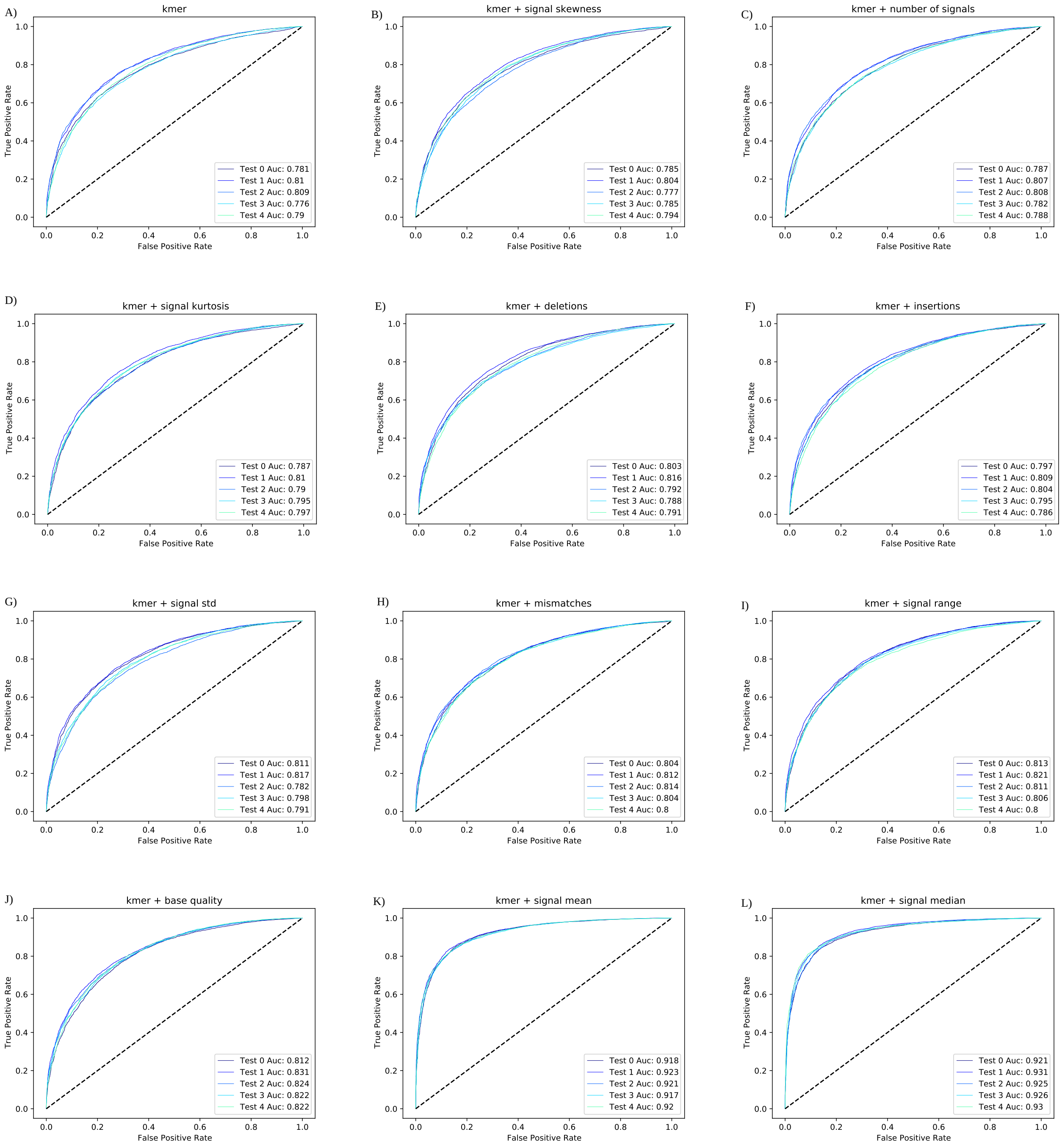


Figure S5: Single feature performance on a subset of the human dataset. Kmer serves as a baseline feature and it is combined with all other individual features. Performance on **A)** kmer alone, **B)** signal skewness, **C)** number of signals, **D)** signal kurtosis, **E)** deletions, **F)** insertions, **G)** signal std, **H)** mismatches, **I)** signal range, **J)** base quality, **K)** signal mean and **L)** signal median. **A-L)** Receiver operating characteristic (ROC) curve showing the false positive rate (x-axis) vs. the true positive rate (y-axis) for the read predictions on a mixture of methylated and unmethylated reads. This analysis was conducted through a 5-fold cross-validation (Test 0-4) using 100,000 labeled examples of human data.

Table S1: Nanopore sequencing datasets employed in the benchmarking and the number of reads corresponding to every dataset

| Species | Name | Reads | Reference |
|---|---|---|---|
| *E. coli* | K12 ER2925 (Total) | 181,137 | Simpson et al. [13] |
| | M.SssI | 69,899 | |
| | Untreated | 111,238 | |
| H. sapiens | NA12878 | | Jain et al. [7] |
| | Norwich | 716,247 | |
| pUC19 | (Total) | 44,511 | Rand et al. [12] |
| | Treated | 17,219 | |
| | Untreated | 27,292 | |

Table S2: Cross validation per-read performance of DeepMP against Nanopolish, DeepSignal, Guppy, Megalodon and DeepMP Seq on *E. coli*.

| Dataset | Test | Method | AUC | Accuracy | Precision | Recall | F-Score |
|---|---|---|---|---|---|---|---|
| *E.coli* | [0, 1000000] | Nanopolish | 0.8783 | 0.7914 | 0.7476 | 0.8834 | 0.8099 |
| | | DeepSignal | 0.9617 | 0.873 | 0.877 | 0.8644 | 0.8707 |
| | | Guppy | 0.9236 | 0.8388 | **0.9962** | 0.654 | 0.7896 |
| | | Megalodon | 0.9804 | 0.9354 | 0.99 | 0.8783 | 0.9308 |
| | | DeepMP Seq | 0.9697 | 0.8687 | 0.8584 | 0.8762 | 0.8672 |
| | | DeepMP | **0.9869** | **0.9368** | 0.9339 | **0.9387** | **0.9363** |
| | [1000000, 2000000] | Nanopolish | 0.8794 | 0.7966 | 0.7621 | 0.8844 | 0.8187 |
| | | DeepSignal | 0.9711 | 0.8943 | 0.9049 | 0.8867 | 0.8958 |
| | | Guppy | 0.9169 | 0.8242 | **0.9961** | 0.636 | 0.7763 |
| | | Megalodon | 0.9789 | 0.9291 | 0.9895 | 0.8709 | 0.9264 |
| | | DeepMP Seq | 0.9685 | 0.882 | 0.9217 | 0.8411 | 0.8795 |
| | | DeepMP | **0.9869** | **0.9372** | 0.9351 | **0.9418** | **0.9389** |
| | [2000000, 3000000] | Nanopolish | 0.877 | 0.7886 | 0.7448 | 0.8873 | 0.8098 |
| | | DeepSignal | 0.9749 | 0.903 | 0.9083 | 0.8989 | 0.9036 |
| | | Guppy | 0.9213 | 0.83106 | **0.9961** | 0.6414 | 0.7803 |
| | | Megalodon | 0.9801 | 0.933 | 0.9901 | 0.8762 | 0.9297 |
| | | DeepMP Seq | 0.9709 | 0.8898 | 0.8971 | 0.8833 | 0.8902 |
| | | DeepMP | **0.9883** | **0.9387** | 0.953 | **0.9243** | **0.9385** |
| | [3000000, 4000000] | Nanopolish | 0.8799 | 0.7939 | 0.7545 | 0.8879 | 0.8158 |
| | | DeepSignal | 0.9754 | 0.9036 | 0.9158 | 0.8939 | 0.9048 |
| | | Guppy | 0.9222 | 0.8303 | **0.9966** | 0.6458 | 0.7838 |
| | | Megalodon | 0.980 | 0.9319 | 0.9906 | 0.8753 | 0.9294 |
| | | DeepMP Seq | 0.9714 | 0.8911 | 0.8996 | 0.8863 | 0.8929 |
| | | DeepMP | **0.9884** | **0.9409** | 0.9477 | **0.9362** | **0.9419** |
| | [4000000, 5000000] | Nanopolish | 0.8777 | 0.7943 | 0.7586 | 0.8867 | 0.8176 |
| | | DeepSignal | 0.9737 | 0.9002 | 0.9116 | 0.8881 | 0.8997 |
| | | Guppy | 0.9237 | 0.8366 | **0.9957** | 0.6557 | 0.7907 |
| | | Megalodon | 0.9808 | 0.9343 | 0.9906 | 0.8781 | 0.9309 |
| | | DeepMP Seq | 0.9699 | 0.8883 | 0.8914 | 0.8864 | 0.8889 |
| | | DeepMP | **0.9879** | **0.9397** | 0.945 | **0.9347** | **0.9398** |

Table S3: Performance summary at read level of DeepMP against Nanopolish, DeepSignal, Guppy, Megalodon and DeepMP Seq on human and pUC19 datasets.

| Dataset | Test | Method | AUC | Accuracy | Precision | Recall | F-Score |
|---|---|---|---|---|---|---|---|
| H. sapiens | Read level | Nanopolish | 0.9284 | 0.908 | 0.9543 | 0.8949 | 0.9236 |
| | | DeepSignal | 0.9629 | 0.9076 | 0.934 | 0.9191 | 0.9255 |
| | | Guppy | 0.8602 | 0.768 | 0.9568 | 0.6565 | 0.7787 |
| | | Megalodon | 0.9394 | 0.8788 | **0.9594** | 0.8418 | 0.8968 |
| | | DeepMP Seq | 0.966 | 0.9146 | 0.9352 | **0.9278** | 0.9315 |
| | | DeepMP | **0.967** | **0.916** | 0.9388 | 0.9260 | **0.9324** |
| pUC19 plasmid | Read level | Nanopolish | 0.7668 | 0.7211 | 0.6038 | 0.7737 | 0.6783 |
| | | DeepSignal | 0.9123 | 0.8141 | 0.8207 | 0.7976 | 0.8090 |
| | | Megalodon | 0.8901 | 0.7730 | 0.6583 | **0.9547** | 0.7792 |
| | | DeepMP Seq | 0.935 | 0.8415 | **0.857** | 0.8108 | 0.8347 |
| | | DeepMP | **0.945** | **0.8697** | 0.8370 | 0.9141 | **0.8738** |

### References

[1] Gpc methylation from fast5's to reference anchored calls - github. github.com/kpalin/gcf52ref. Retrieved June 2021.

[2] Live methylation calling blog. simpsonlab.github.io/2020/03/03/ nanopolish-v0.12.0/. Retrieved June 2021.

[3] Oxford nanopore technologies. community.nanoporetech.com/. Retrieved June 2021.

[4] Oxford nanopore technologies github - megalodon. github.com/nanoporetech/megalodon. Retrieved June 2021.

[5] Quickstart - calling methylation with nanopolish. nanopolish.readthedocs.io/en/latest/. Retrieved June 2021.

[6] Rerio models - github. github.com/nanoporetech/rerio. Retrieved June 2021.

[14] Marcus Stoiber, Joshua Quick, Rob Egan, Ji Eun Lee, Susan Celniker, Robert K Neely, Nicholas Loman, Len A Pennacchio, and James Brown. De novo identification of dna modifications enabled by genome-guided nanopore signal processing. *BioRxiv*, page 094672, 2016.

[15] Zaka Wing-Sze Yuen, Akanksha Srivastava, Runa Daniel, Dennis McNevin, Cameron Jack, and Eduardo Eyras. Systematic benchmarking of tools for cpg methylation detection from nanopore sequencing. *Nature Communications*, 12(1):1–12, 2021.
